## Supplementary Information for "The brain hierarchically represents the past and future during multistep anticipation"

### Story Examples

“I was relaxing in the meadows one day when I came upon an interesting hallway; I decide to walk through. When I go through the last door of the hallway, I land on this futuristic platform that tells me to pick a letter on the screen. I pick the third letter, and get transported into a huge, dark, scary cave. I frantically run to the end of the cave, and it leads me to a beautiful sunny park. At the park, I am asked to pick between the yellow gazebo and the red gazebo. I pick the yellow one, and I find myself in a tunnel. Once I go through the archway of this tunnel, I realize that it’s a portal. I zoom through the portal, and end up in my house on Christmas day of 2005. I realize that I've time travelled. After enjoying a meal with my family, I enter the fireplace and am transported back to the meadow where I began.”

“I am being held captive by an intergalactic military convoy. I am being taken to a prison on a remote forest world that is covered in darkness every day. I am dragged along a dark path by the brutish guards, and after a night of travel they leave me in this cell. What it presents as upscale boutique aesthetics festers into cold minimalism and rots the brain of the inhabitant. But I escape. A cleverly timed throwing-a-chair-through-the-window leaves me stranded in the barren hills that make up this planet. I find myself staggering across each hill, thinking each vista might hold sight of my salvation. One of them does, in the form of an old pub with a blinking neon sign and a game of dominoes going on outside. I order a drink. A man, dressed in a monocle and festive top hat turns to me, asks me my story. I tell it. He seems amused. He invites me to his summer palace on a brighter part of the planet, that is only reachable through hot air balloon.

It is warm. I can feel the suns tapping warmly on my cheek. The eccentric man puts his hands on my shoulders. You will love it there, he says. When we arrive, I can see why he might have thought so. But why come all this way to take a mere stranger back with you. It has been days. It is beautiful on his estate. He never takes off his festive tophat, not even over eggs in the morning or with a nightcap over the fire. After a week he invites me into his study, in the wing he had banned me from entering. Upon seeing the walls, it washes over me in an instant. This is the leader of the Christmas empire, he knew of my escape and took me captive. He laughs.

Guards enter the room and take me aboard the ship that will take me to prison. This is the last time I let my guard down for a festive top hat."

### Simulation of BOLD Data

We conducted simulations of BOLD data to investigate whether variation in signal-to-noise and $R^2$ was confounded with the width of the Gaussian fit across regions. To investigate this, we systematically simulated datasets by adding varying amounts of noise to visual cortex data and assessing changes in the resulting Gaussian fits for each dataset.

We added random noise to the visual cortex patterns of activity elicited during the cue screen of the Anticipation Task and to the across-participant environment templates elicited during the Localizer Task (random values drawn from a normal distribution centered at 0). To create datasets with different amounts of noise, we varied the standard deviation of this normal distribution from 0.1 to 2 in increments of 0.1. We then ran the same Gaussian analysis as reported in the manuscript on each of these simulated datasets. We simulated five datasets at each noise level.

The amplitude of the Gaussian fits became successively smaller in datasets with successively more noise (Supplementary Fig 4a and 4d). Additionally, $R^2$ became successively smaller in datasets with successively more noise (Supplementary Fig 4b). Critically, however, the forward and backward sigmas did not become wider in datasets with successively more noise (Supplementary Fig 4c and 4d). Instead, as amplitude and $R^2$ decreased, the forward and backward sigmas remained similar. Therefore, our simulations reveal that (1) decreased signal to noise ratio does not explain the variation in Gaussian widths and (2) variation in goodness of fit does not account for changes in Gaussian width, as $R^2$ and width vary independently in our simulations.

### Supplementary Figures

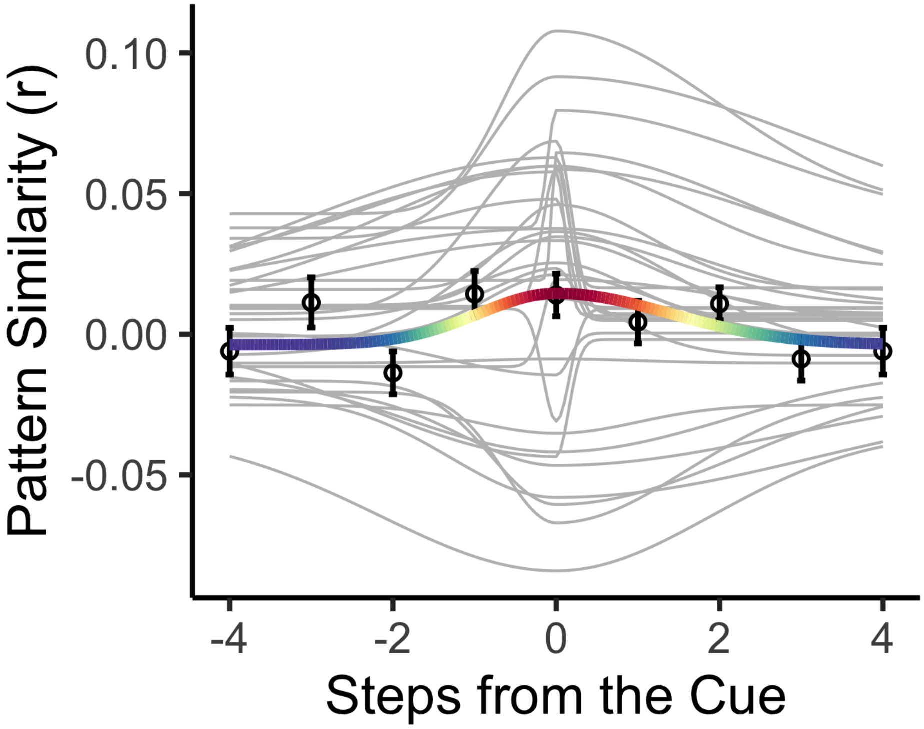


**Supplementary Figure 1 | Gaussian Curve in Insula.** The Gaussian curve in the insula was not a significantly better fit for the correctly ordered data vs. the shuffled null including the cue, indicating a lack of temporal structure representations in this region. Points indicate average pattern similarity at each step from the cue and error bars indicate standard error of the mean. Thick, colored line indicates the average Gaussian fit across participants, with the red end of the rainbow scale indicating higher pattern similarity and the purple end indicating lower pattern similarity. Thin gray lines indicate participant-specific Gaussian fits. N = 32 participants.

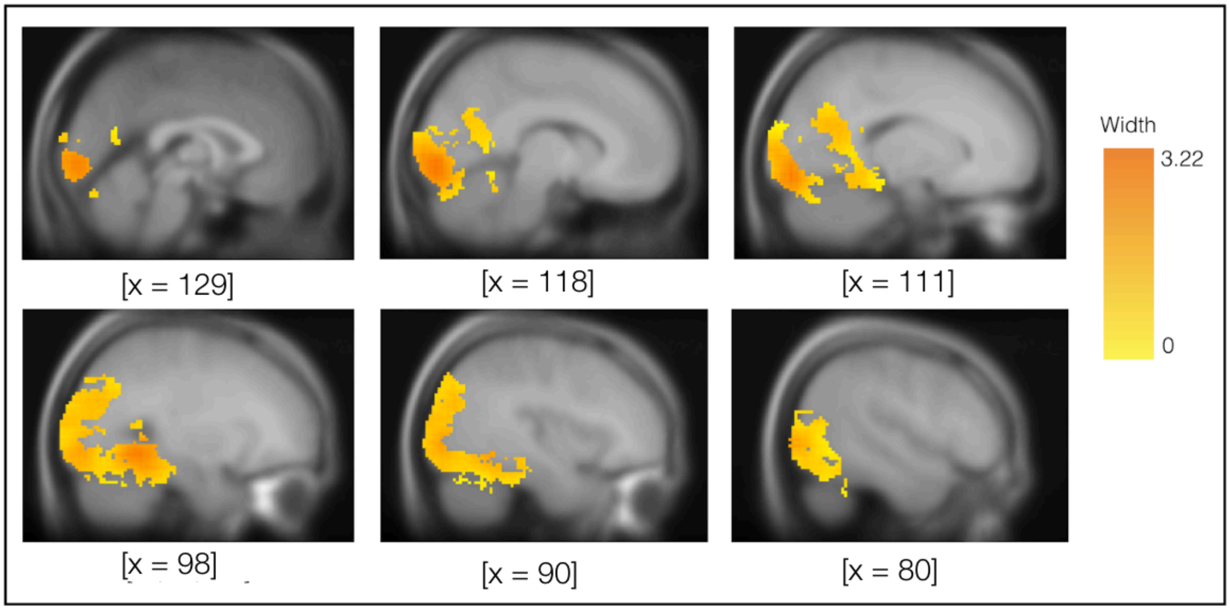


**Supplementary Figure 2 | Searchlight results.** Voxels with significant Gaussian fits were found across the visual system. Voxels are colored based on the average of their Gaussians' forward and backward widths. MNI coordinates are indicated in square brackets.

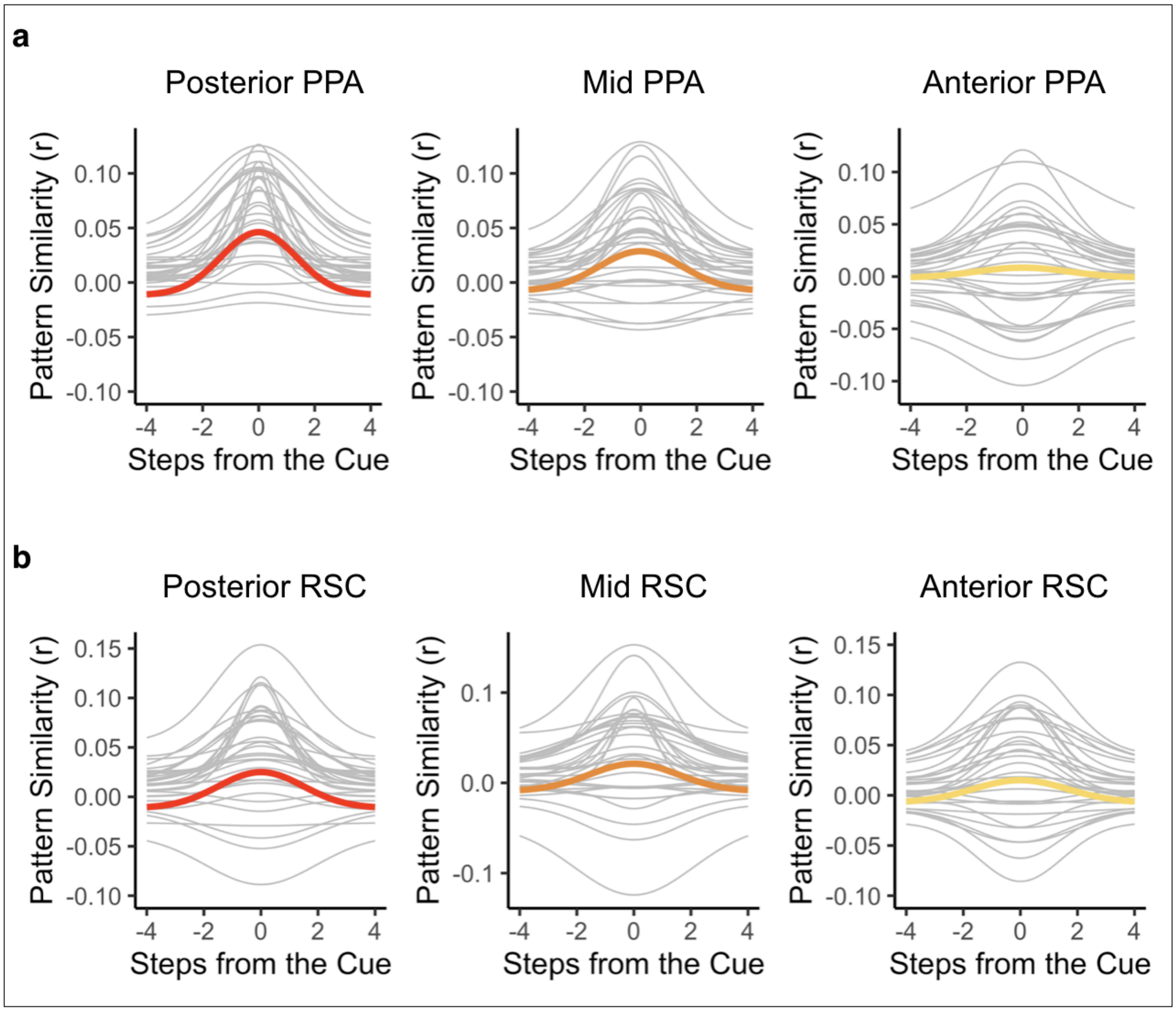


**Supplementary Figure 3 | Participant-specific Gaussian Curves of Sample Voxels in PPA and RSC.** Gaussian fit from an exploratory searchlight in PPA **(a)** and RSC **(b)** for three randomly selected voxels in posterior (left, red), mid (middle, orange), and anterior (right, yellow) parts of both regions. Gray lines indicate participant-specific Gaussian curves and solid colored lines indicate the average Gaussian fit across participants. N = 32 participants.

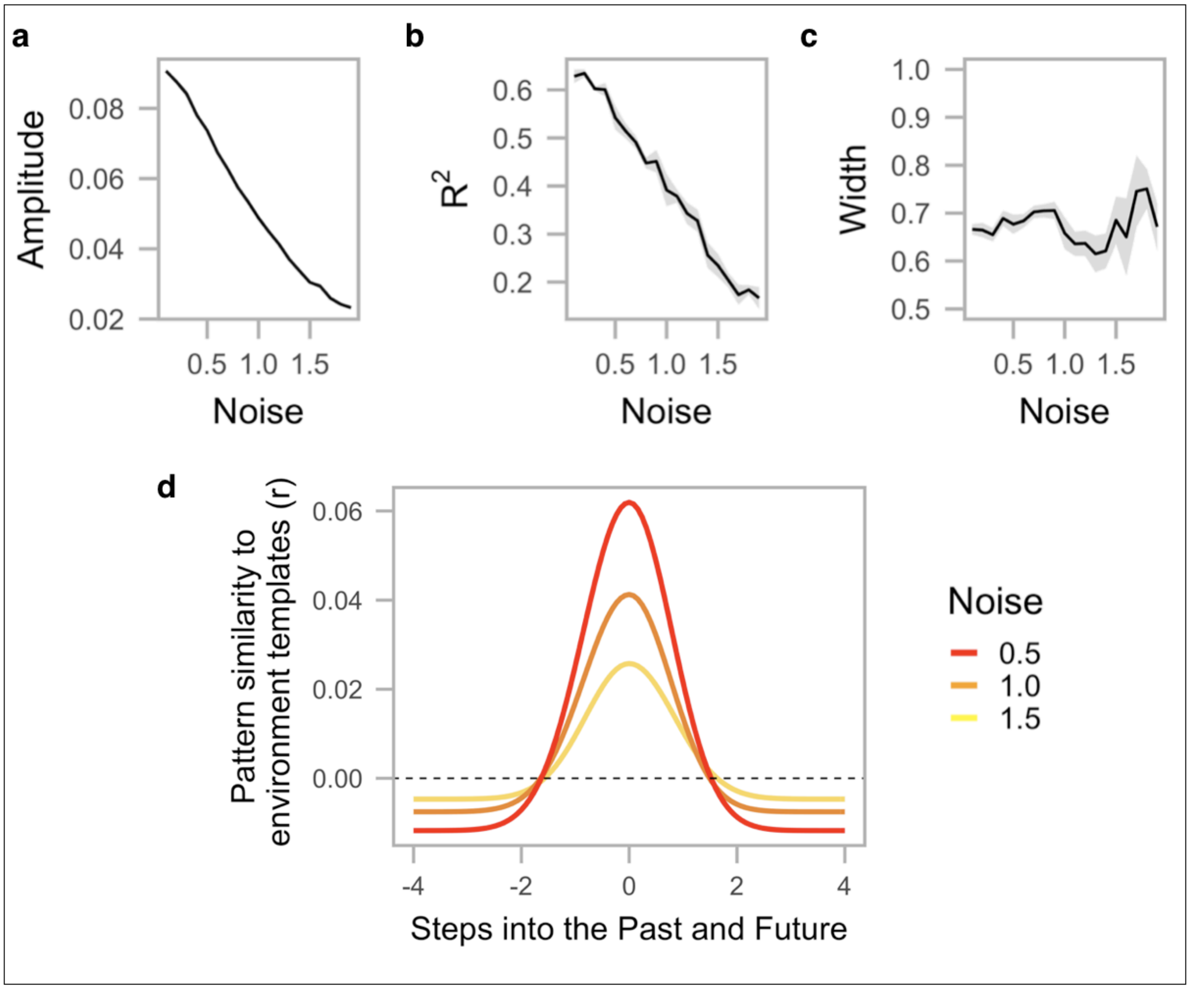


**Supplementary Figure 4 | Results of BOLD Data Simulation.** The amplitude **(a)** and $R^2$ **(b)** of the Gaussian fit decreased for simulated datasets with successively higher noise levels, but there was no systematic change in the width of the Gaussian fit **(c)**. Black lines indicate the average value across five simulated datasets at each noise level and transparent gray ribbons indicate the 95% confidence interval. Note that the error ribbon is also shown in panel **(a)** but difficult to detect due to very low variability. **(d)** Gaussian curve at three sample noise levels.

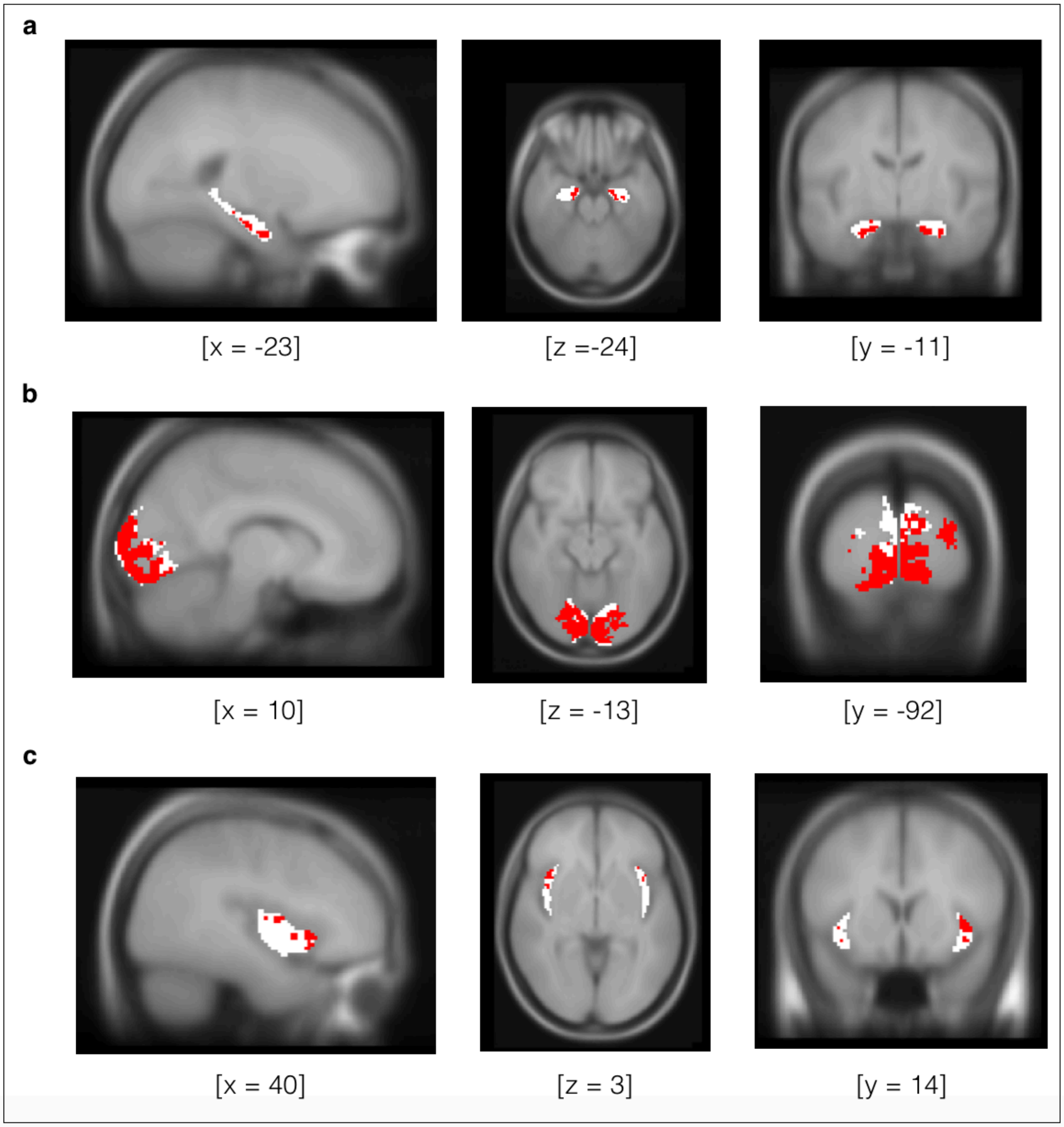


**Supplementary Figure 5 | Regions of Interest.** Visualization of the three *a priori* ROIs analyzed in the experiment. Anatomical ROIs for hippocampus **(a)**, visual cortex **(b)**, and insula **(c)** are shown in white. Within each anatomical ROI, voxels that reliably discriminated between environments during the Localizer Task are shown in red. The voxels in red make up the conjunction ROIs used in our analyses. MNI coordinates are indicated in square brackets.

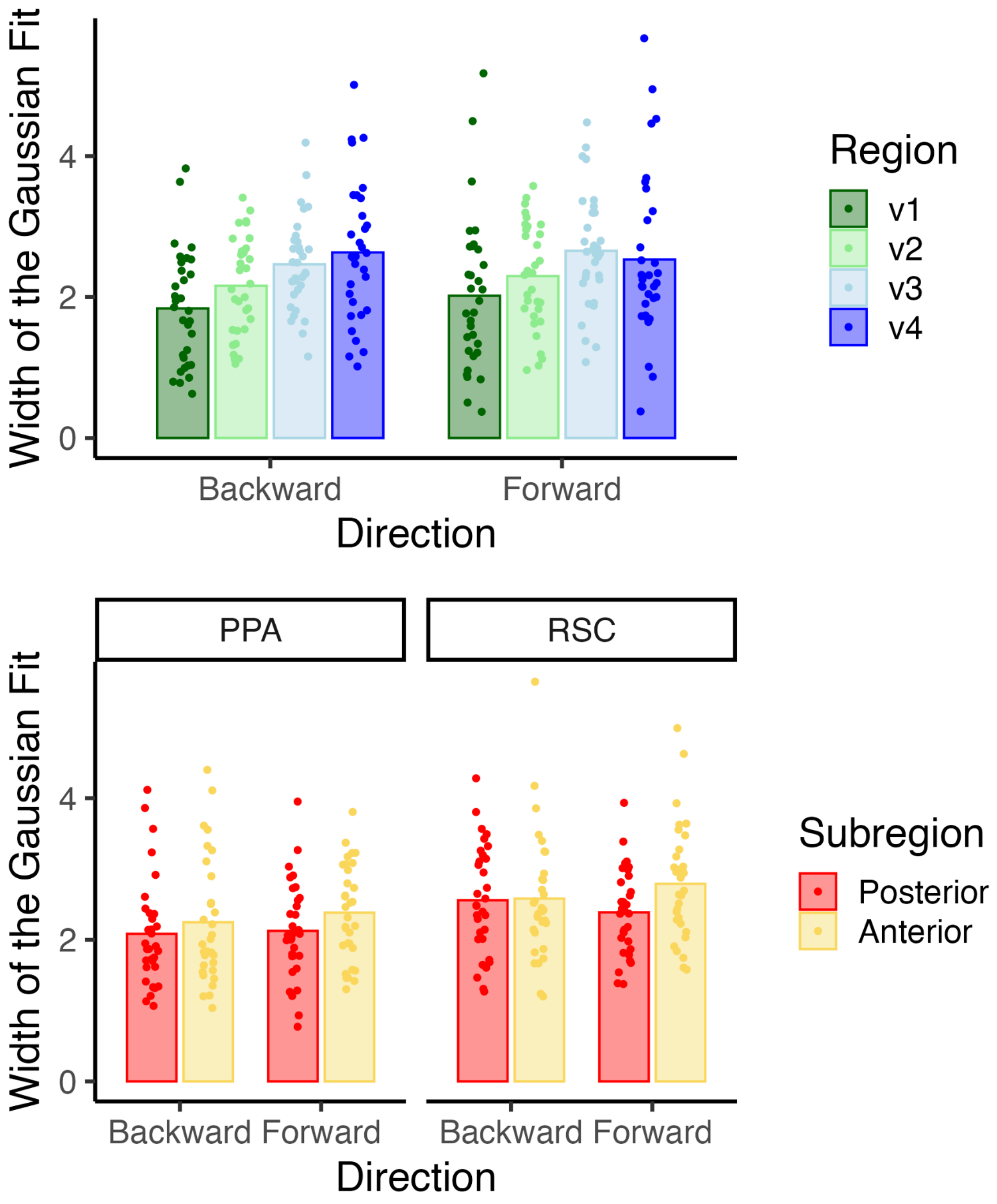


**Supplementary Figure 6 | Forward and Backward Widths Across Visual Regions.** The backward and forward width of the Gaussian fit increased hierarchically across early visual cortex from V1 to V4 (Top). In PPA, the forward and backward width of the Gaussian fit increased from posterior to anterior voxels. In RSC, the forward, but not the backward, width of the Gaussian fit increased from posterior to anterior voxels (Bottom). Data are shown binned by anterior vs. posterior aspects of PPA and RSC, but the statistics reported in the main text treated the anterior-posterior axis of these regions as a continuous variable. Bars indicate the average width across participants and points indicate the width of the Gaussian fit for each participant. N = 32 participants.
